# Highly Potent Peptide Therapeutics To Prevent Protein Aggregation In Huntington’s Disease

**DOI:** 10.1101/2023.04.27.538016

**Authors:** Anooshay Khan, Esra Yuca, Cemile Elif Özçelik, Ozge Begli, Oguzhan Oguz, Serkan Kasırga, Urartu Ozgur Safak Seker

**Affiliations:** UNAM-Institute of Materials Science and Nanotechnology, Bilkent University, 06800, Ankara, Turkey; Department of Neurosciences, Bilkent University, Ankara, Turkey; Department of Molecular Biology and Genetics, Yildiz Technical University, Istanbul, Turkey; Interdisciplinary Neuroscience Program, Bilkent University, 06800 Ankara, Turkey

**Keywords:** Huntington’s Disease, Peptide-based Drug therapy, Huntingtin, Htt aggregation, inhibition

## Abstract

Huntington’s disease (HD) is a progressive, autosomal dominant neurodegenerative disorder resulting from a significant amplification of CAG repeats in exon 1 of the Huntingtin (Htt) gene. More than 36 CAG repeats result in the formation of mutant Htt (mHtt) protein. These amino-terminal mHtt fragments lead to the formation of misfolded proteins, which then form aggregates in relevant brain regions. Available treatments concentrate primarily on alleviating the disease’s symptoms. Therefore, therapies that can delay the progression of the disease are imperative to halt the course of the disease. Peptide-based drug therapies provide such a platform. Inhibitory peptides were screened against monomeric units of both wild type (Htt(Q25)) and mHtt fragments, including Htt(Q46)and Htt(Q103). It was accomplished by utilizing several display technologies. This study focuses on the in-vitro characterization of the screened peptides. Fibril kinetics was studied in real-time utilizing the Thioflavin T (ThT) assay. The impact of specific peptides on fibril formation was examined by observing the change in fluorescence signal. Atomic force microscopy was also used to study the influence of peptides on fibril formation. Three of the six chosen peptides (HHGANSLSLVSQD, HGLHSMHNKLTR, and WMFPSLKLLDYH) effectively inhibited aggregation. These experiments demonstrate that the chosen peptides suppress the formation of fibrils in mHtt proteins and can provide a therapeutic lead for further optimization and development.

## 1. INTRODUCTION

Huntington’s disease (HD) is a progressive autosomal-dominant neurological disorder that causes cognitive decline, behavioral disturbances, and neuropsychiatric symptoms ^1^. HD is caused by a mutation in exon1 of the Htt gene, located on chromosome 4 (4p16.3). The gain of function mutation causes dramatic expansion of glutamine encoding CAG sequences that exceed 35 repeats, causing misfolded protein. Typically, individuals with the clinical phenotype have 27 to 35 repeats in their genome. It is commonly believed that HD can develop when 36 repeats or more are present, although full penetration does not occur until 40 ^2^. High CAG repeats count has also been linked to earlier onset, faster progression, and increased severity of the disease ^3^.

This CAG-PolyQ expansion results in misfolded protein, which is highly prone to aggregation. The aggregated mutant Htt (mHtt) protein forms inclusion bodies inside the cells, leading to cell death and neurodegeneration ^4^. mHtt aggregation process has three stages: the initial lag phase and an exponential growth phase, followed by a plateau (Figure 1). Primary nucleation initiates aggregate formation in mHtt by templated aggregation ^5^. The N-terminal domain of exon1 of mHtt is flanked by polyQ repeats, and conformational changes in this region accelerate nucleation. The polyQ repeats form β-sheets which aggregate by hydrogen bonds. The aggregates then form protofibrils and elongate into fibrils by adding new monomers as they grow. Fibril-dependent nucleation is observed in mHtt, resulting in a branched morphology unlike other amyloid fibrils (Amyloid Beta, Alpha-synuclein) ^6^. It is claimed that protein-protein interactions, post-translational changes, and protein cleavage all contribute to the pathogenesis of HD ^7^. HD is a rare, age-onset degenerative disease with a prevalence of 2.7 per 100,000 worldwide and has been a subject of scientific and medical fascination for decades ^8^.

**Figure 1.**
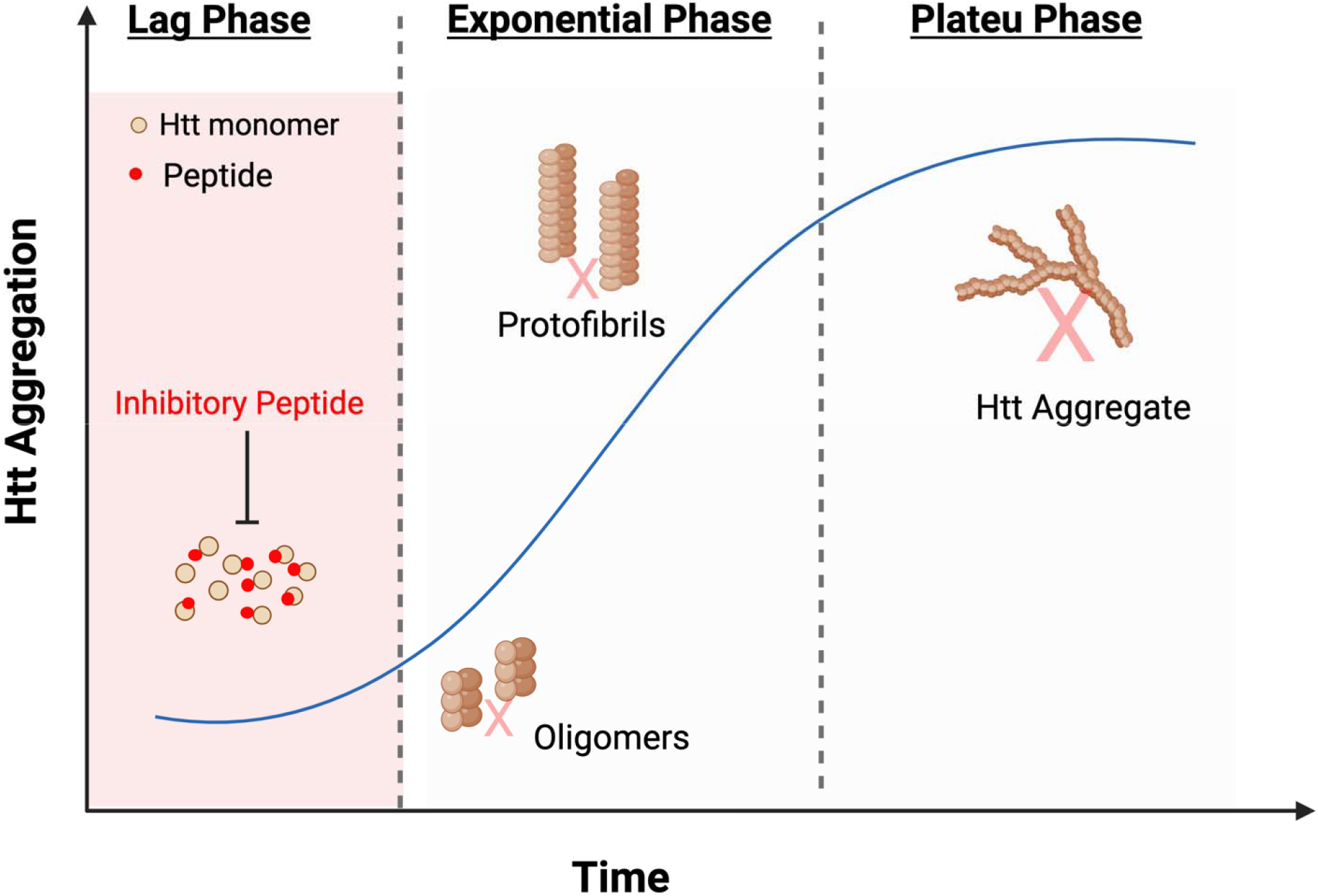
The effect of the proposed inhibitory peptides on the aggregation kinetics of Htt proteins. The inhibitory peptides bind to the monomeric units of the Htt proteins, preventing aggregate formation.

There is currently no viable treatment for HD that reduces the disease’s severity and progression. Available pharmacological treatments for HD are inadequate and mainly focus on alleviating the symptoms of the disease rather than halting its development. In response to these deficiencies, small molecules (ASO ^9^, RNAi ^9^, antibody ^10^) and peptide-based therapeutics ^11^ are being investigated. Peptides offer several benefits over small compounds, including high specificity, strong biological activity, cheap cost, and excellent membrane penetrability ^11^.

Peptide-based therapeutics for HD have shown promising results by hampering disease progression by preventing aggregation. Some of the research emphasized in this respect includes the following: polyQ region (25Q) separated by an alpha-helix colocalized and interacted with polyQ Htt protein aggregates; this peptide was able to minimize and delay aggregate development ^12^. Polyglutamine binding peptide 1 (QBP1) and bivalent Htt-binding peptide were found to be particularly effective in binding abnormal polyQ proteins ^12, 13^. However, these peptides mainly focus on inhibiting aggregation later in the cascade of Htt polymerization and aggregate development.

The candidate inhibitory peptides tested in this study were screened against the monomeric units of Htt proteins, both wild-type (Htt(Q25)) and mHtt fragments such as Htt(Q46) and Htt(Q103) ^14^. The monomeric units of these proteins were expressed on the yeast surface, which allowed for proper eukaryotic folding/misfolding. The peptides were introduced by the phage display library. The ligand peptides were selected by biopanning without immobilizing yeast cells ^14^. The capacity of peptide to bind monomeric Htt fragments permits the inhibition of aggregation at a very early stage in the course of disease progression. It lends originality to the putative peptides employed in this investigation. This study focuses on the in vitro characterization of ligand peptides that have been previously screened. In this investigation, we examined whether the screened peptides inhibited the fibril formation of mHtt proteins. Aggregation kinetics of the mHtt proteins with and without peptide treatment was investigated by ThT assay and Atomic Force Microscopy. As a result, three out of six proposed peptides significantly blocked the aggregation of Htt proteins.

## 2. MATERIALS AND METHODS

### 2.1. Expression and Purification of Htt Constructs

GST tag was fused to stabilize and prevent the polymerization of Htt proteins, as described previously ^15^. Both wildtype GST-Htt-(Q25) and mutant GST-Htt-(Q46) and GST-Htt-(Q103) fusion proteins were cloned in pET22b (+) vector under the T7 promoter. A TEV protease cleavage site was cloned between the 6His-GST and N terminal of Htt Exon1 proteins. The plasmids harboring the cloned constructs were transformed in *E. coli* BL21 (DE3) for protein expression. The details of the cloning, expression and purification of all the GST-Htt constructs used in this study is given in the Supporting Information (Figures S1−S3 and Table S1-S2).

### 2.2. ThT Assay to Visualize Aggregation in Real-time

The peptides used in this study were produced by solid-state synthesis and were purchased from Peptiteam. For initiating aggregation, TEV protease was added to 7 μM of GST-Htt protein. 7 μM of each peptide was then added to the peptide-treated groups. The reaction was completed to 200 μL in each well by tev reaction buffer, and 0.5 μL of 25 μM ThT was added. The 96-microwell plate was then sealed with adhesive tape to prevent evaporation. The ThT fluorescence intensity of each sample was recorded by spectraMax M5 Microplate Reader spectrophotometer (Molecular Devices) with 438/495 nm excitation/emission filters at cutoff at 475 nm after every 10 minutes for 16-20 hours. Each reading was preceded by 5 s for orbital agitation. All readings were set up as triplicates. The ThT data were normalized as described before ^16^, and percentage fluorescence was plotted against time (Figure S5).

### 2.3. Aggregation Kinetics

Freshly purified Htt proteins were centrifuged at 18000 x g for 20 minutes to remove pre-existing aggregates ^17^. The TEV cleavage reaction was set up to initiate aggregation using 10 μM GST-Htt(Q103). 500 μL volumes of reactions were incubated at room temperature. 2 hours after the administration of TEV protease, 10 μM peptide was introduced at time zero. After 2,4,8,10,12 and 24 hours, 5 μL of samples were aliquoted from both peptide-treated and untreated Htt(Q103) for AFM deposition. On SiO_2_ wafers, a 5 μL sample was deposited in order to get topography pictures in air. The wafer was cleaned with ultrapure water for two minutes and then dried with argon.

In order to examine the impact of inhibitory peptides on the aggregation kinetics of GST-Htt(Q46) and GST-Htt(Q25), the purified proteins were diluted to a final concentration of 30 μM in 1X TEV reaction buffer (pH 8.0) and TEV Protease was added to remove 6xHis-GST and commence the aggregation. 500 μL volumes of reactions were incubated at room temperature. 2 hours after the administration of TEV protease, equimolar peptide was introduced at time zero.

After 4,8, and 24 hours, 5 μL of samples were aliquoted from the peptide-treated and untreated Htt(Q25) and Htt(Q46) experimental groups. The sample deposition for AFM was carried out in the same manner.

### 2.4. AFM Imaging

AFM images of AFM surface topography were captured with an Oxford Instruments MFP-3D Origin in tapping mode, which employs air topography. Budget AFM probes with a 300 kHz resonance frequency and 40 N/m spring constant Sensors were used for measuring purposes.

Samples were scanned between 1.2Hz and 2.2Hz, depending on the size of the scan area. AFM measurements were conducted in ambient conditions at room temperature. Matlab FiberApp was used to quantify each data set’s fibril length. Matlab AFM pictures were evaluated and adjusted for scan artifacts using the Gwyddion v2.60 application. The two-way ANOVA test compared the fibril length of peptide-treated Htt proteins and untreated Htt proteins.

## 3. RESULTS

### 3.1. The Effect of Inhibitory Peptides on Protein Aggregation

To determine the effect of inhibitory peptides on protein aggregation in real time, we performed ThT assay. Initiating the aggregation with the cleavage of the GST solubility tag guarantees monomeric Htt starting conditions, which were otherwise challenging to accomplish as the protein polymerizes as soon as the tag is removed. We started ThT measurements soon after the addition of TEV protease. By adding an equimolar inhibitory peptide to the protein, the effect of the peptide on the fibrilization and consequent aggregation was evaluated. As the length of polyQ repeats correlates directly with the acceleration of aggregation kinetics, tht assay revealed that Htt(Q103) exhibited the highest fluorescence, followed by Htt (Q46) and Htt(Q25) (Figure 2b). Fluorescence was monitored for a minimum of 16 hours at 37 °C.

**Figure 2.**
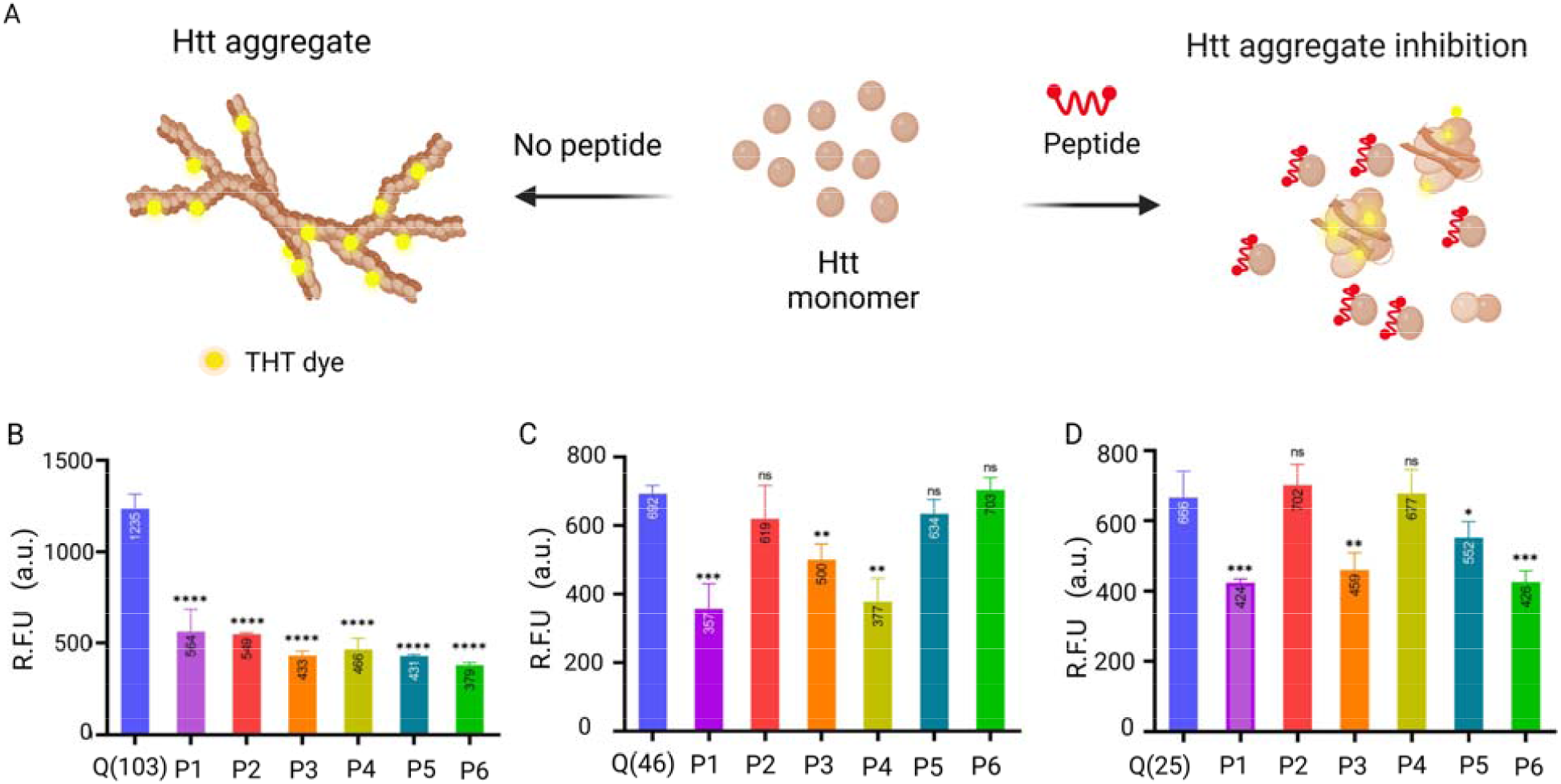
ThT assay to evaluate aggregation kinetics of peptide treated and control Htt proteins. (A) Shows ThT dye binding to the beta-sheet structure of Htt fibrils. In the presence of peptides that inhibit the Htt fibril elongation, the binding of ThT dye is reduced. (B-D) ThT fluorescenc intensity was recorded for Htt protein (B) Htt(103Q) (C) Htt(46Q) and (D) Htt(Q25) in the presence of peptides (1-6) at a 1:1 molar ratio. ThT emission was recorded at 495 nm upon excitation at 438 nm. The bar graph averages the endpoint reading at 16 hours over triplicates for each sample. (P<0.0001)

We knew from the previous study that these candidate inhibitory peptides interact with the monomeric units of Htt proteins ^14^. Here we wanted to evaluate whether the interaction enabled or blocked the aggregation in the Htt proteins. We saw that all six peptides decreased fluorescence significantly in Htt(Q103) (Figure 2). The fluorescence values were normalized, and the graphs were plotted (Figure S5). The decrease in fluorescence indicates that the peptide hindered the fibrillization process and that the inhibitory peptide successfully prevented protein aggregation. When studying fibril kinetics with Htt(Q46) with the peptides, we saw that only peptide1, peptide 3, and peptide 4 decreased the fluorescence signal significantly (Figure 2B). Peptide 5 and peptide 2 also decreased the fluorescence signal; however, the decrease was insignificant according to the student-t-test. Peptide 6, on the other hand, increased the fluorescence signal, indicating that the addition of peptide promoted aggregation in Htt(Q46) protein.

Unlike mutant Htt exon 1 fragments, wild-type Htt with less than 36 polyQ repeats does not form insoluble but soluble aggregates ^18^. Baseline fluorescence was observed for Htt(25Q), attributed to the presence of beta sheets (although of shorter length and with fewer cross-links). Since the screened peptides were screened against all three Htt (Q25), (Q46), and (Q103) proteins, the impact of the candidate inhibitory peptides on wild-type Htt (Q25) was also investigated. Peptide1, 3 and 5, and 6 showed a decrease in fluorescence signal for Htt Q25, and peptides 2 and 4 led to increased fluorescence (Figure 2B), hence increasing aggregation. After quantifying the fluorescence signals and comparing the effects of peptides on fibril kinetics, shown in table S3, we shortlisted three peptides that caused a decrease in aggregation in all three proteins tested. These peptides were peptide 1, peptide 3, and peptide 5. When comparing various proteins and peptides, the intensity of fluorescence by Tht assay is typically a insufficient quantitative indicator of fibril production and inhibition. Therefore, we wanted to test further shortlisted peptides’ effect on aggregation kinetics by Atomic Force microscopy.

### 3.2. Morphological Characteristics of Htt Nanofibrils

We examined the morphological characteristics of huntingtin nanofibrils at various phases of in-vitro fibril formation, both in the presence and absence of peptides. As done previously, the aggregation process was commenced by cleaving GST with site-specific TEV protease. After allowing the fibrils to grow for two hours, peptides were added. This time was recorded as t=0 hours. As seen in Figure 3(D), for both control and peptide-added proteins, spherical oligomers and individual fibrils were observed for Htt(Q103), and only spherical oligomers were observed for Htt(Q46) and Htt(Q25) at t=0. As previously reported, these should be noticed during the first phases of fibrillization kinetics ^17^. Samples for AFM imaging were taken every 4 hours (Figures S6-S8). At t=4, we noticed an increase in the length and branching of fibrils. Branched morphology is a well-established property of polyQ > 36 Htt proteins, and this was seen for t=4, t=8, and t=24 for both Htt(Q103) and Htt(Q46). Over time, the branching increased, and clusters of aggregates were observed, except for Htt(Q25). At each time point, the length of 100 distinct fibrils was measured. The length of fibrils at various time points is represented graphically in Figure 3. At t=24, a reduction in length was noticed. This reduction may result from the protein degradation in the TEV cleavage buffer (pH 8.0).

**Figure 3.**
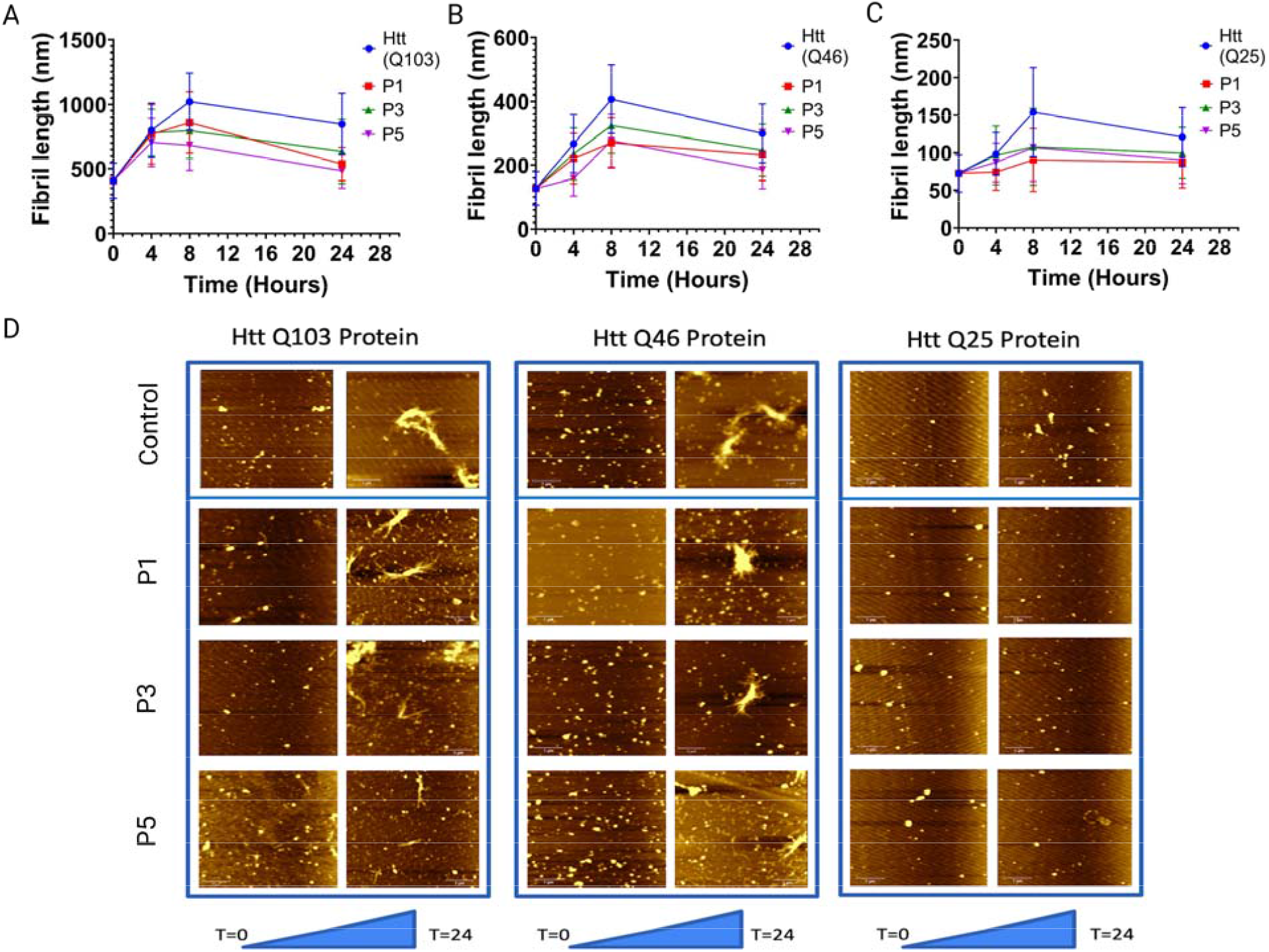
Kinetics of aggregation of the Htt(Q103), Htt(Q46) and Htt(Q25) protein. (A) Shows change in fibril length both control Htt(Q103) and peptide treated Htt(Q103) with respect to time. (B) Shows change in fibril length both control Htt(Q46) and peptide treated Htt(Q46) with respect to time. (C) Shows change in fibril length both control Htt(Q25) and peptide treated Htt(Q25) with respect to time. The fibril length was measured after 0, 4, 8, and 24 hours and the mean length of 100 fibrils and standard deviation at each time point is plotted. (D) Shows AFM images of all three proteins both in the presence and absence of peptides, taken at t=0 and t=24 of the aggregation. All images show large-scale views with Scale bars are 1 μm.

### 3.3. Kinetics of Aggregation of The Htt(Q103), Htt(Q46) and Htt(Q25) Protein

Comparing the fibril kinetics of peptide-treated Htt(Q103), a substantial reduction in fibril length was seen at all time points. 24 hours after peptide addition, we observed that peptide 1 led to a 36.7% decrease in fibril length, peptide 3 caused a 22.67% decrease, and peptide 5 caused a 28% decrease in the fibril length. In addition, a decline in clustering and branching was also observed. This reduction was assessed by measuring 100 fibrils at each time point and graphing the average lengths (Figure 3). Even at early stages of aggregation kinetic, peptide inhibits agglomeration. At t=4 and t=8, less branching and even isolated fibrils are visible. The reduction in fibril length at an early stage of fibril production confirms that the peptides interact with the monomeric units of the htt proteins. This data provides conclusive proof that the applied peptide inhibits the aggregation of the ordinarily aggregated Htt(Q103) protein, which exhibits distinct branching and clustering. Similarly, the fibril kinetics of peptide-treated Htt(Q46) also showed a reduction in fibril length at all time points. 24 hours after peptide treatment, we observed that peptide 1 led to a 24.91% decrease in fibril length, peptide 3 led to a 17.61% length decrease, and peptide 5 caused a 17.3% decrease in fibril length.

While studying aggregation kinetics for Htt(Q25), tiny circular oligomers with a diameter of 80-100 nm were seen for Htt(Q25) (protein alone) at t=0. The dimensions of the circular oligomers were in accordance with what was predicted. Since Htt(Q25) does not form insoluble aggregates, most t=4 samples consisted of small oligomers with relatively few individual fibrils. These fibrils were considerably shorter than the mutant Htt exon1 proteins mentioned before. At t=8, protein-only tiny individual fibrils measuring 100-140 nm were found for Htt(Q25), and at t=24, small oligomers were again observed. These oligomers’ diameters were measured. 24 hours after peptide treatment, we observed that peptide 1 led to a 42.87% decrease in fibril length, peptide 3 led to a 38.33% length decrease, and peptide 5 caused a 25.6% decrease in the fibril length. In order to remove any contaminant in the SiO_2_ wafer, only oligomers taller than 5nm were considered. The sizes of 100 distinct fibrils/oligomers were measured at each time point.

The length of these fibrils at various time intervals is shown in Figure 3 (C). At time t=24, a reduction in length and protein degradation were detected.

By comparing the fibril dynamics of peptide-treated proteins, a substantial reduction in fibril length was seen at all time points. No individual fibers were seen at periods 4, 8, and 24; only oligomers were observed for all peptides. Even for short poly-Q stretches, our data demonstrate that the inhibitory peptides are highly selective and block fibril formation.

## 4. DISCUSSION

Overall results of the AFM and the ThT assay complement each other. It provides solid evidence that the inhibitory peptides effectively inhibited the in-vitro aggregation of wild-type and mutant Htt fibrils. Exon1 of both mutant and wild-type htt was the focus of this research. Previous research has demonstrated that N-terminal Htt peptides exhibit high pathogenicity ^19^. The expansion of the poly-Q stretch in exon1 causes specific structural and physicochemical alterations in Htt proteins, resulting in misfolded monomers that quickly polymerize into hazardous oligomers. These oligomers subsequently form inclusion bodies in the brain’s basal ganglia, impairing neuronal function and resulting in neural degeneration ^20^. Since HD is an autosomal dominant condition, wild-type Htt was included in all the aggregation kinetics investigations as one mutant allele is sufficient for pathological aggregation formation.

Furthermore, it has been demonstrated that mutant Htt(Q46) seeds may induce the formation of insoluble aggregates on Htt(Q25) in vitro ^21^. Therefore, we want to determine whether or not our peptides affected the generation of soluble aggregates by Htt(Q25) and whether or not our peptide treatment will be beneficial in heterozygous HD variants. Using peptides that prevent this aggregation in Htt protein by interacting with polyQ stretches provides a promising platform for drug development. These peptides’ low potential toxicity and ability to target the mHtt at the initial steps in the aggregation make them ideal candidates for peptide-based therapeutics.

The inhibitory peptides tested in this study were screened against the monomeric units of Htt proteins by display technologies.

The ability of the peptide to bind monomeric htt fragments allows the blocking of aggregation at a very early stage of disease progression. This approach gives novelty to our proposed peptides used in this study.

Future studies will focus on assessing the toxicity of these selected peptides in mammalian cells; both would do this in vitro and in vivo experiments.

In vitro studies will utilize the HT22 (mouse hippocampal) cell line. Furthermore, the toxicity will be evaluated by MTT (3–(4,5-dimethylthiazol-2-yl)-2,5-diphenyltetrazolium bromide) colorimetric test for viability assay. The peptides will be tested individually on Ht22 cells and in combination with Htt proteins in an equimolar ratio by co-transfection. This will allow us to evaluate two things; whether The peptide has any toxic effect on the neurons, and whether the peptide is decreasing the toxicity of the mHtt aggregates in the neurons. After this, the peptides that pose the least to no toxicity will be selected for in vivo experiments in animal models.

## 5. CONCLUSION

In this study, candidate ligand peptides were characterized and evaluated for their inhibitory effect against three Htt proteins; (Q25), (Q46) and (Q103). In vitro characterization of peptides on aggregation, kinetics was studied by ThT Assay and AFM. 3 of the 6 selected peptides (HHGANSLSLVSQD), (HGLHSMHNKLTR) and (WMFPSLKLLDYH) showed a decrease in fluorescence signal, indicating inhibition in fibril formation for all three proteins. AFM images for these peptides showed less aggregation in mHtt proteins than untreated mHtt proteins. These studies show that the selected peptides are effective for inhibiting the aggregation of fibrils in mHtt proteins and are promising candidates for peptide therapy for HD.

## GRAPHICAL ABSTRACT

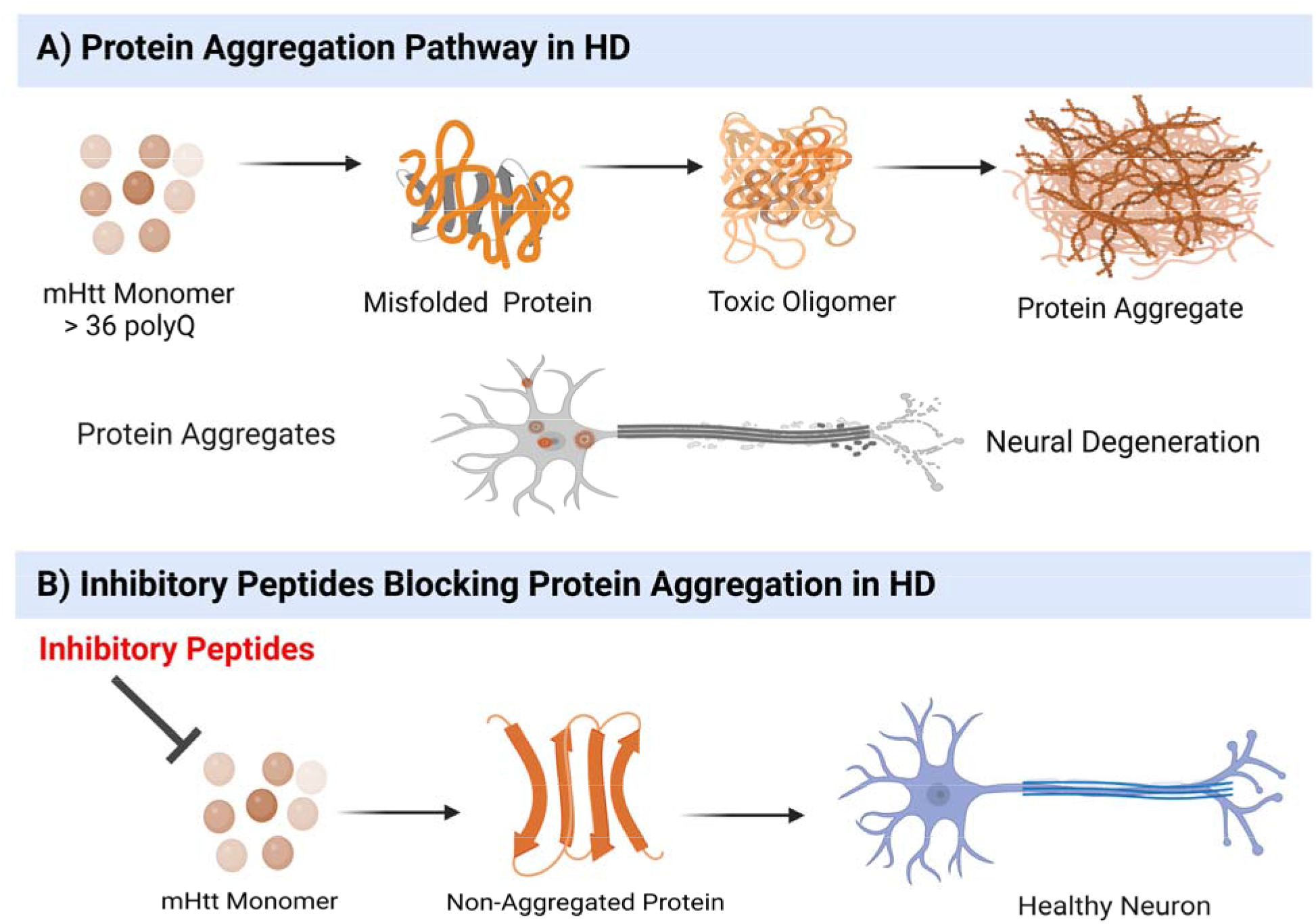

## ASSOCIATED CONTENT

### Supporting Information

Additional experimental details, materials, and methods, including photographs of experimental setup (DOC)

## AUTHOR INFORMATION

## Author Contributions

UOSS conceived the idea. UOSS, AK and EY planned the experiments for molecular cloning, protein expression and purification. AK, EY and CEO, OB carried out experiments. OO and SK carried out AFM imaging experiments.

## ACKNOWLEDGMENT

We thank Recep Erdem Ahan, and Ahmet Hincer for their support with the experiments.

## ABBREVIATIONS

Htt: Huntingtin
ThT: Thiofavin T
AFM: Atomic Force Microscopy

