## Supplementary document for "Highly Potent Peptide Therapeutics To Prevent Protein Aggregation In Huntington’s Disease"

Supplementary Data

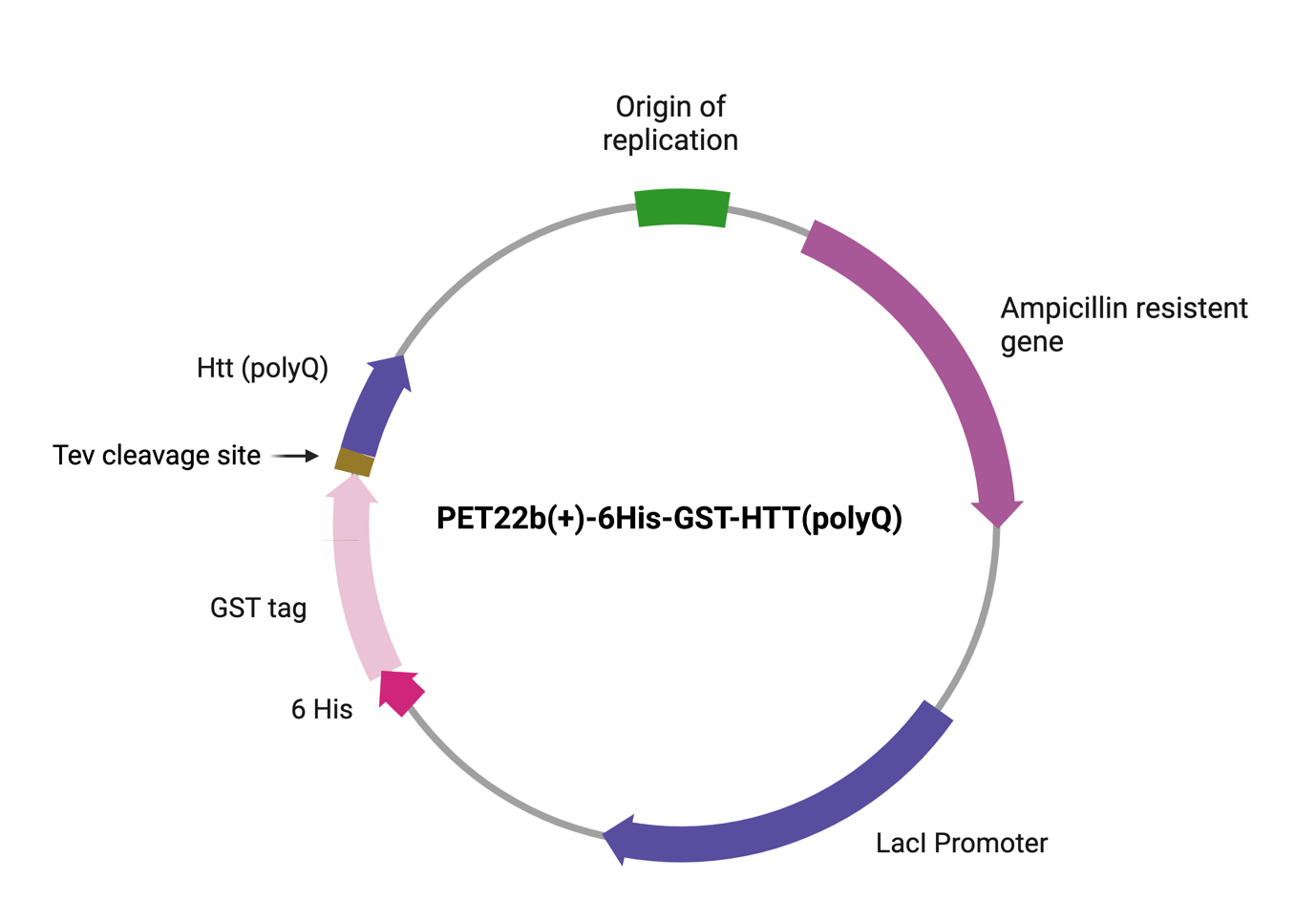

Figure S1.Plasmid map of PET22b(+) vector with GST-Htt construct.

| Htt constructs | | Sequence |
| --- | --- | --- |
| GST-Htt(Q25) | ATGAACACCATTCATCACCATCACCATCACAACACTAGTATGTCCCCTATACTAGGTTATTGGAAAATTAAGGGCCTTGTGCAACCCACTCGACTTCTTTTGGAATATCTTGAAGAAAAATATGAAGAGCATTTGTATGAGCGCGATGAAGGTGATAAATGGCGAAACAAAAAGTTTGAATTGGGTTTGGAGTTTCCCAATCTTCCTTATTATATTGATGGTGATGTTAAATTAACACAGTCTATGGCCATCATACGTTATATAGCTGACAAGCACAACATGTTGGGTGGTTGTCCAAAAGAGCGTGCAGAGATTTCAATGCTTGAAGGAGCGGTTTTGGATATTAGATACGGTGTTTCGAGAATTGCATATAGTAAAGACTTTGAAACTCTCAAAGTTGATTTTCTTAGCAAGCTACCTGAAATGCTGAAAATGTTCGAAGATCGTTTATGTCATAAAACATATTTAAATGGTGATCATGTAACCCATCCTGACTTCATGTTGTATGACGCTCTTGATGTTGTTTTATACATGGACCCAATGTGCCTGGATGCGTTCCCAAAATTAGTTTGTTTTAAAAAACGTATTGAAGCTATCCCACAAATTGATAAGTACTTGAAATCCAGCAAGTATATAGCATGGCCTTTGCAGGGCTGGCAAGCCACGTTTGGTGGTGGCGACCATCCTCCAAAAGGATCCGAAAACCTGTATTTTCAGGGCactagtATGGCGACCCTGGAAAAGCTGATGAAGGCCTTCGAGTCCCTCAAAAGCTTCCAACAGCAGCAACAGCAACAACAGCAGCAACAGCAACAACAGCAGCAACAGCAACAACAGCAGCAACAGCAACAACCGCCACCACCTCCCCCTCCACCCCCACCTCCTCAACTTCCTCAACCTCCTCCACAGGCACAGCCTCTGCTGCCTCAGCCACAACCTCCTCCACCTCCACCTCCACCTCCTCCAGGCCCAGCTGTGGCTGAGGAGCCTCTGCACCGACCT | |
| GST-Htt(Q46) | ATGAACACCATTCATCACCATCACCATCACAACACTAGTATGTCCCCTATACTAGGTTATTGGAAAATTAAGGGCCTTGTGCAACCCACTCGACTTCTTTTGGAATATCTTGAAGAAAAATATGAAGAGCATTTGTATGAGCGCGATGAAGGTGATAAATGGCGAAACAAAAAGTTTGAATTGGGTTTGGAGTTTCCCAATCTTCCTTATTATATTGATGGTGATGTTAAATTAACACAGTCTATGGCCATCATACGTTATATAGCTGACAAGCACAACATGTTGGGTGGTTGTCCAAAAGAGCGTGCAGAGATTTCAATGCTTGAAGGAGCGGTTTTGGATATTAGATACGGTGTTTCGAGAATTGCATATAGTAAAGACTTTGAAACTCTCAAAGTTGATTTTCTTAGCAAGCTACCTGAAATGCTGAAAATGTTCGAAGATCGTTTATGTCATAAAACATATTTAAATGGTGATCATGTAACCCATCCTGACTTCATGTTGTATGACGCTCTTGATGTTGTTTTATACATGGACCCAATGTGCCTGGATGCGTTCCCAAAATTAGTTTGTTTTAAAAAACGTATTGAAGCTATCCCACAAATTGATAAGTACTTGAAATCCAGCAAGTATATAGCATGGCCTTTGCAGGGCTGGCAAGCCACGTTTGGTGGTGGCGACCATCCTCCAAAAGGATCCGAAAACCTGTATTTTCAGGGCactagtATGGCGACCCTGGAAAAGCTGATGAAGGCCTTCGAGTCCCTCAAAAGCTTCCAACAGCAGCAACAGCAACAACAGCAGCAACAGCAACAACAGCAGCAACAGCAACAACAGCAGCAACAGCAGCAACAGCAACAACAGCAGCAACAGCAACAACAGCAGCAACAGCAACAACAGCAGCAACAGCAACAACCGCCACCACCTCCCCCTCCACCCCCACCTCCTCAACTTCCTCAACCTCCTCCACAGGCACAGCCTCTGCTGCCTCAGCCACAACCTCCTCCACCTCCACCTCCACCTCCTCCAGGCCCAGCTGTGGCTGAGGAGCCTCTGCACCGACCT | |
| GST-Htt(Q103) | ATGAACACCATTCATCACCATCACCATCACAACACTAGTATGTCCCCTATACTAGGTTATTGGAAAATTAAGGGCCTTGTGCAACCCACTCGACTTCTTTTGGAATATCTTGAAGAAAAATATGAAGAGCATTTGTATGAGCGCGATGAAGGTGATAAATGGCGAAACAAAAAGTTTGAATTGGGTTTGGAGTTTCCCAATCTTCCTTATTATATTGATGGTGATGTTAAATTAACACAGTCTATGGCCATCATACGTTATATAGCTGACAAGCACAACATGTTGGGTGGTTGTCCAAAAGAGCGTGCAGAGATTTCAATGCTTGAAGGAGCGGTTTTGGATATTAGATACGGTGTTTCGAGAATTGCATATAGTAAAGACTTTGAAACTCTCAAAGTTGATTTTCTTAGCAAGCTACCTGAAATGCTGAAAATGTTCGAAGATCGTTTATGTCATAAAACATATTTAAATGGTGATCATGTAACCCATCCTGACTTCATGTTGTATGACGCTCTTGATGTTGTTTTATACATGGACCCAATGTGCCTGGATGCGTTCCCAAAATTAGTTTGTTTTAAAAAACGTATTGAAGCTATCCCACAAATTGATAAGTACTTGAAATCCAGCAAGTATATAGCATGGCCTTTGCAGGGCTGGCAAGCCACGTTTGGTGGTGGCGACCATCCTCCAAAAGGATCCGAAAACCTGTATTTTCAGGGCactagtATGAAGGCCTTCGAGTCCCTCAAAAGCTTCCAACAGCAGCAACAGCAACAACAGCAGCAACAGCAACAACAGCAGCAACAGCAACAACAGCAGCAACAGCAACAACAGCAGCAACAGCAACAACAGCAGCAACAGCAACAACAGCAGCAACAGCAACAACAGCAGCAACAGCAACAACAGCAGCAACAGCAACAACAGCAGCAACAGCAACAACAGCAGCAACAGCAACAACAGCAGCAACAGCAACAACAGCAGCAACAGCAACAACAGCAGCAACAGCAACAACAGCAGCAACAGCAACAACAGCAGCAACAGCAACAACCGCCACCACCTCCCCCTCCACCCCCACCTCCTCAACTTCCTCAACCTCCTCCACAGGCACAGCCTCTGCTGCCTCAGCCACAACCTCCTCCACCTCCACCTCCACCTCCTCCAGGCCCAGCTGTGGCTGAGGAG | |

Tabe Sl : GST-Htt sequences used in this study

| Primers | | Sequence 5’-3' |
| --- | --- | --- |
| pCEO69 | GCAGCCGGATCTCAGTGGTGGTGGTGGTGGTGCTCGAGTCATCAAGGTCGGTGCAGAGGCTC | |
| pCEO-70 | GTTTGGTGGTGGCGACCATCCTCCAAAAGGATCCGAAAACCTGTATTTTCAGGGCactagtATGGCGACCCTGG | |

Table S2. List of primers used to clone the Htt constructs in PET22b(+) AmpR Vector

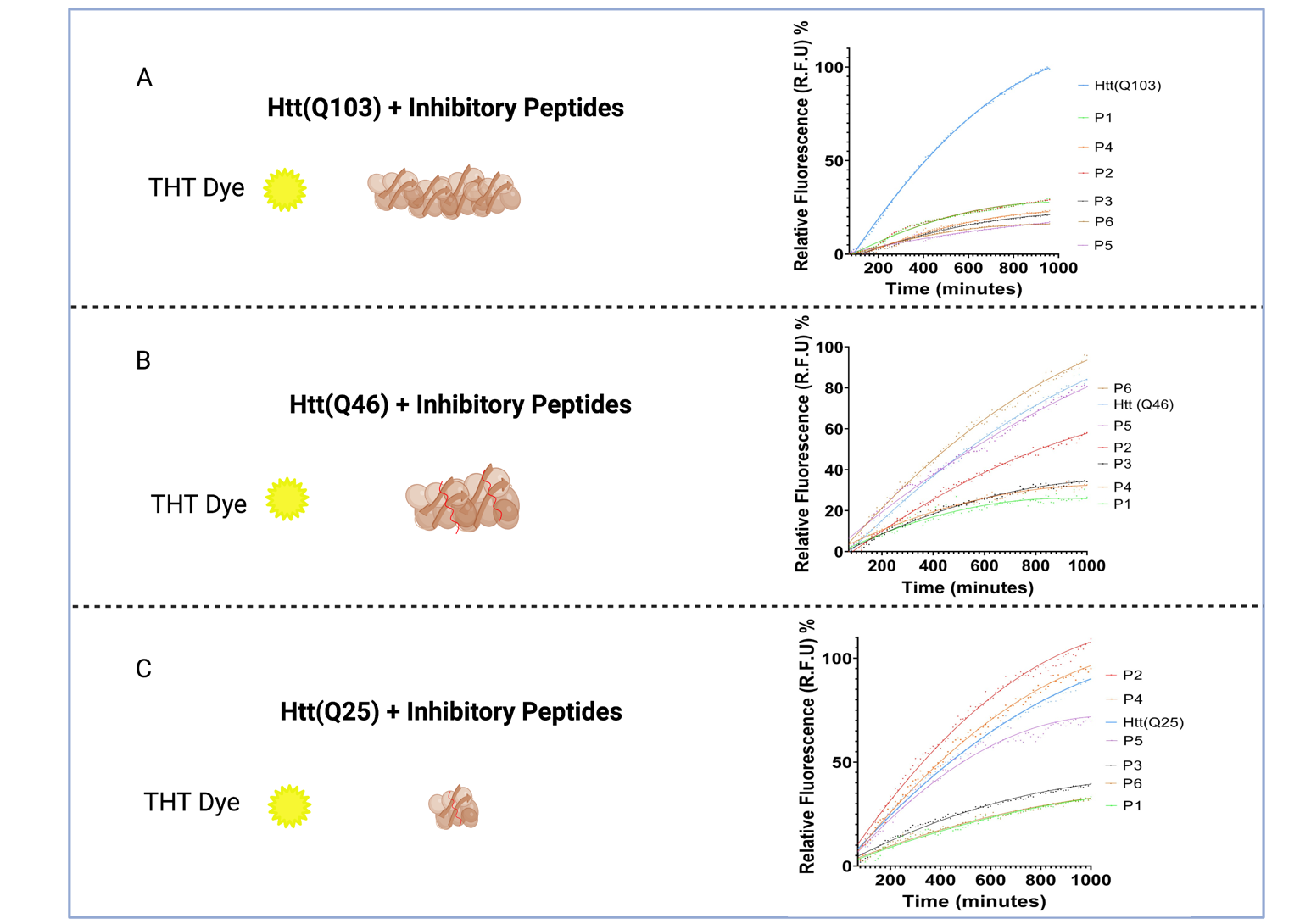

Figure S2. Tht assay for evaluating the effect of peptides on the Fibril kinetics of Htt proteins.

AFM supplementary data

Htt 103 Protein

Protein

P1

P3

P5

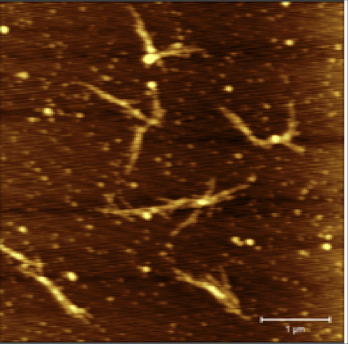

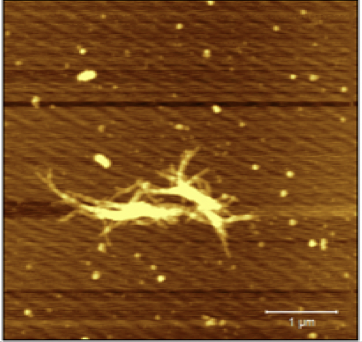

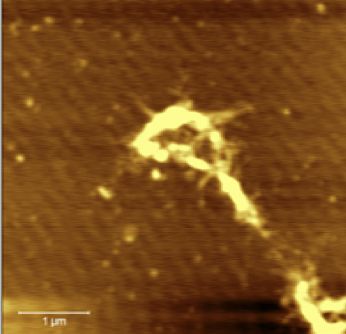

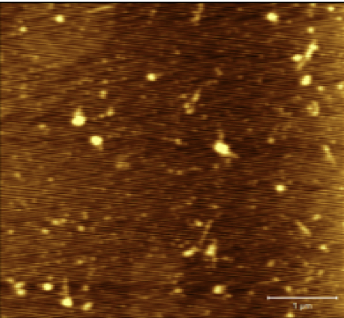

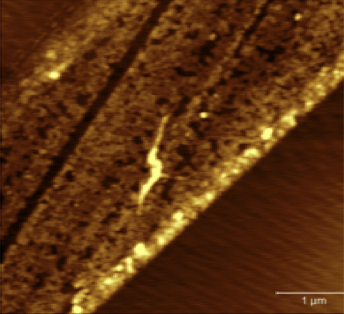

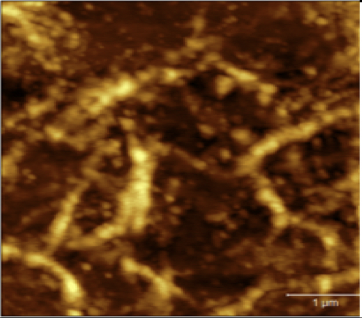

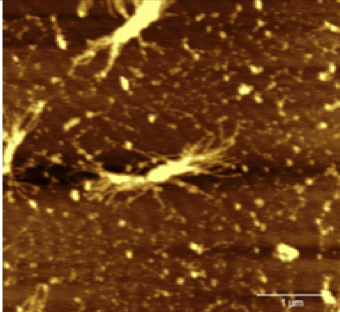

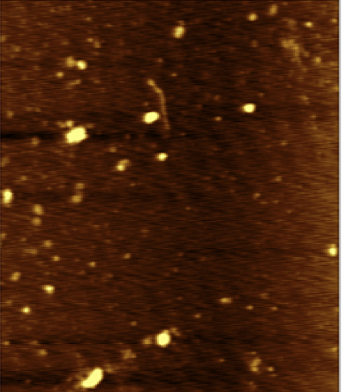

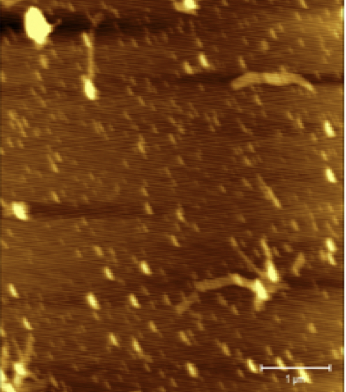

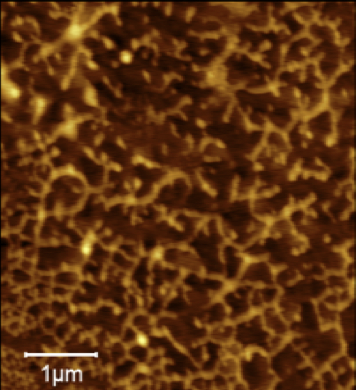

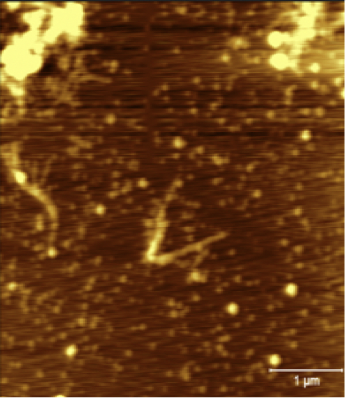

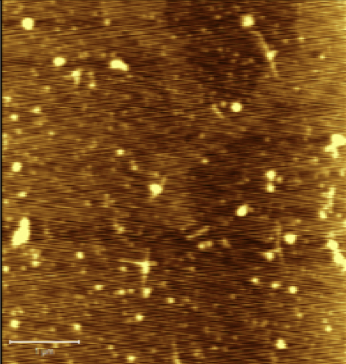

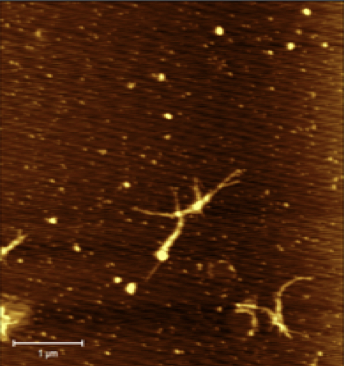

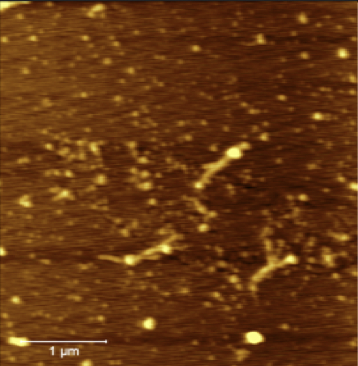

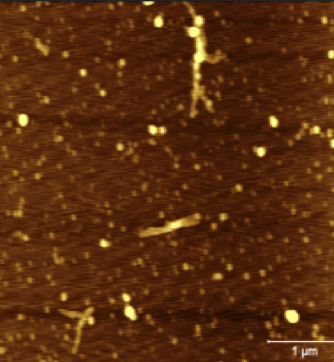

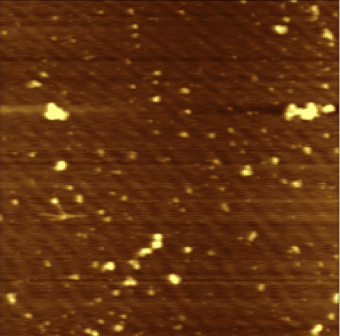

T= 4

T=0

T=8

T=24

Figure S3. Shows the aggregation kinetics of HttQ103, both in the absence and presence of inhibitory peptides. The AFM images show samples obtained at 0 hour, 4 hours, 8 hours, and 24 hours. The first row of Images corresponds to the fibril growth kinetic of the Htt Q103 protein. The second row of images corresponds to the fibril growth kinetics of HttQ103 in the presence of inhibitory peptide 1 (**HHGANSLLGLVQS).** The third row shows fibril growth kinetics of HttQ103 in the presence of inhibitory peptide 3 (**HGLHSMHNKLLQT)** and the last row shows fibril growth kinetics of HttQ103 in the presence of inhibitory peptide 5 (**WMFPSLKLLDYH). Scale bar is 1**

Htt (Q46) Protein

Protein

P1

P3

P5

T= 4

T=0

T=8

T=24

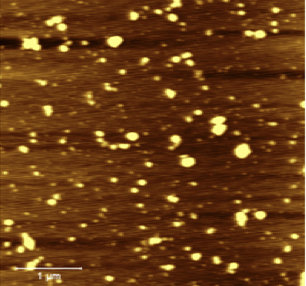

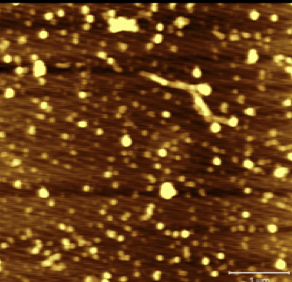

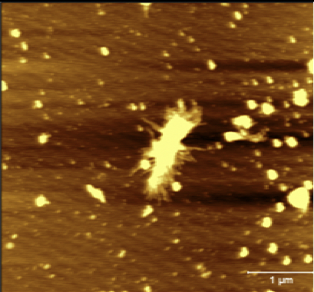

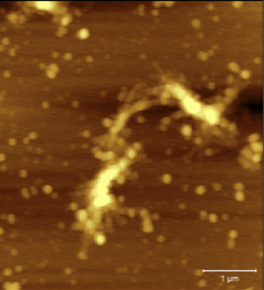

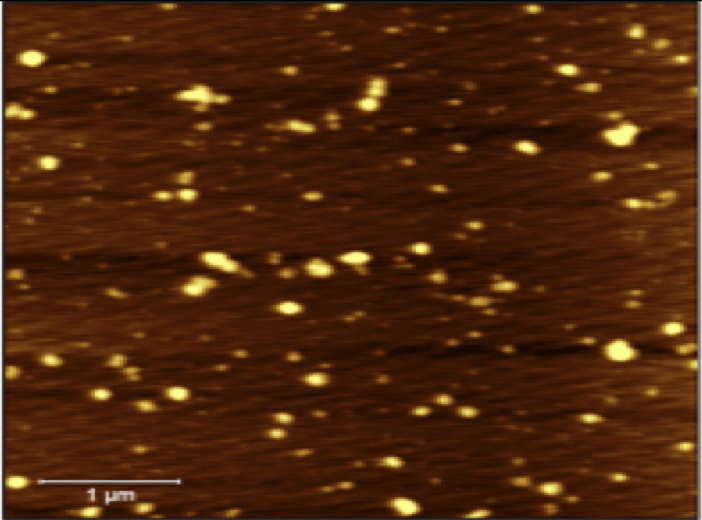

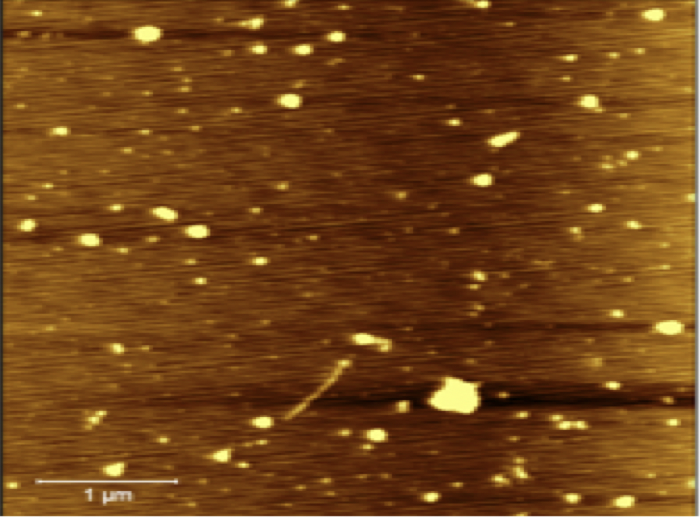

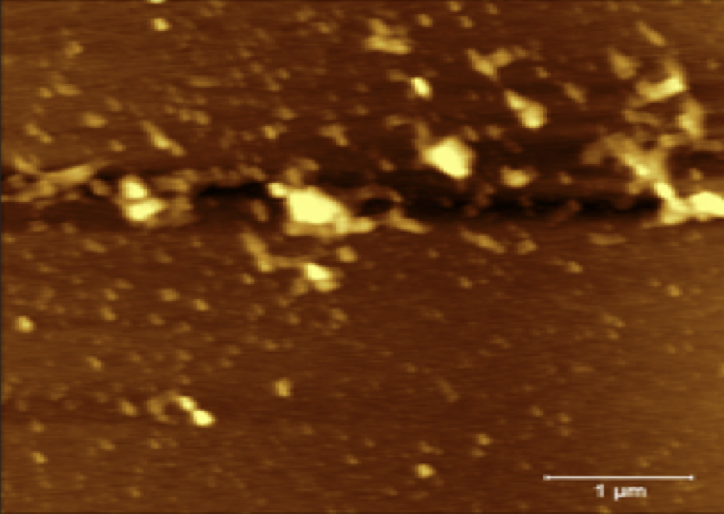

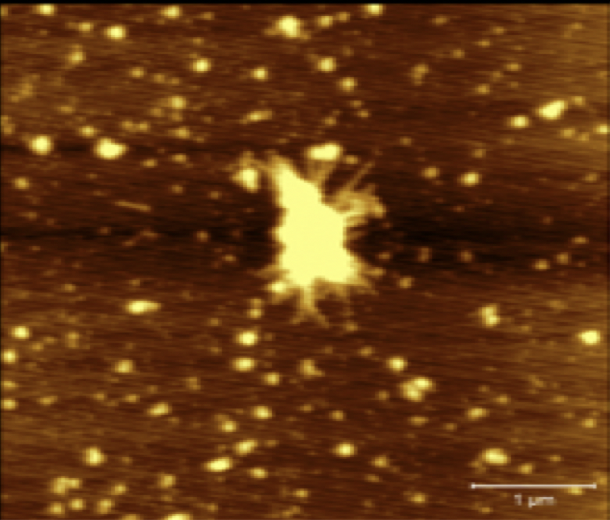

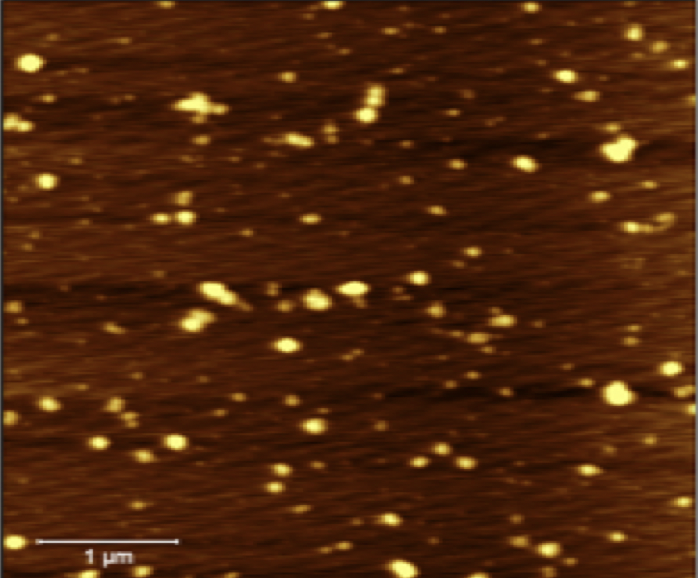

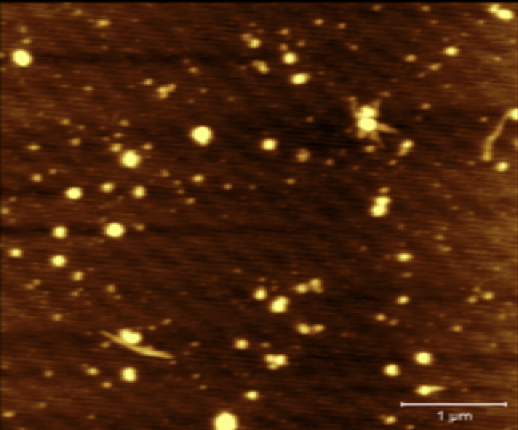

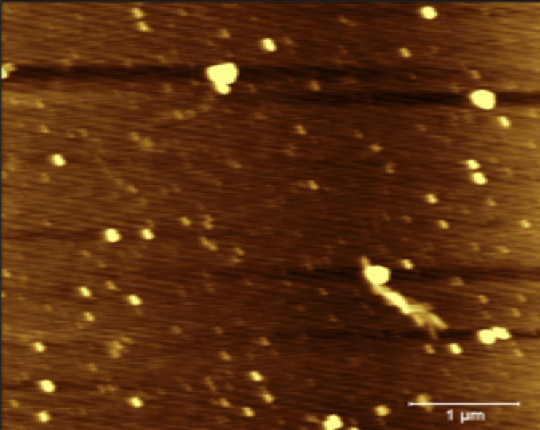

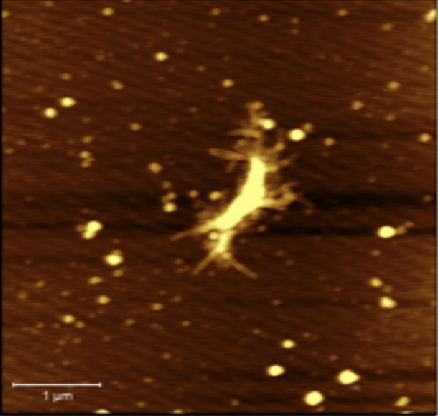

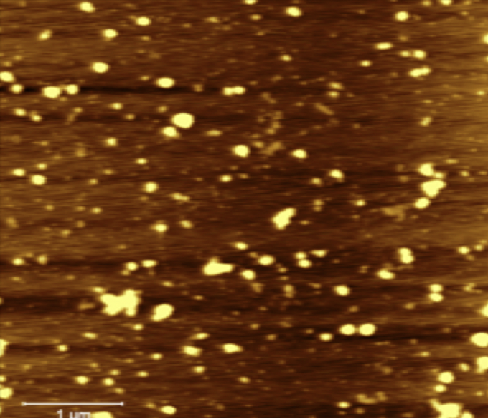

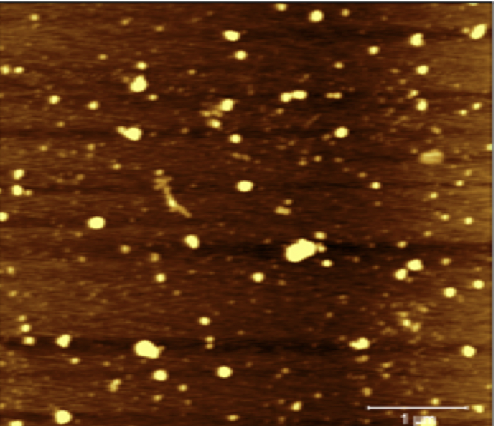

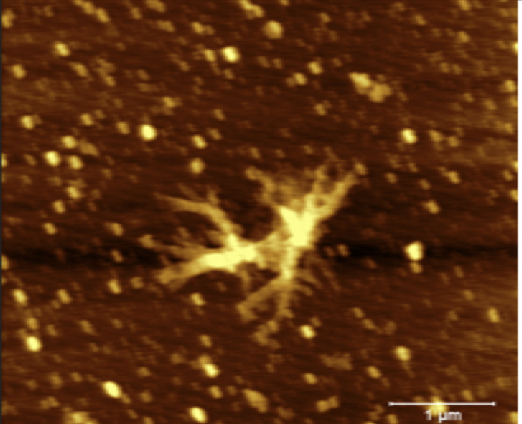

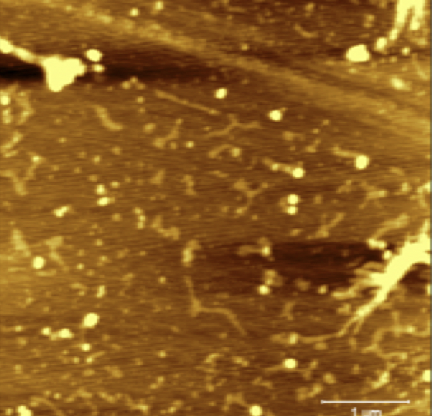

Figure S4. Shows the aggregation kinetics of HttQ46, both in the absence and presence of inhibitory peptides. The AFM images show samples obtained at 0 hour, 4 hours, 8 hours, and 24 hours. The first row of Images corresponds to the fibril growth kinetic of the Htt Q46 protein. The second row of images corresponds to the fibril growth kinetics of HttQ46 in the presence of inhibitory peptide 1 (**HHGANSLLGLVQS).** The third row shows fibril growth kinetics of HttQ46 in the presence of inhibitory peptide 3 (**HGLHSMHNKLLQT)** and the last row shows fibril growth kinetics of HttQ46 in the presence of inhibitory peptide 5 (**WMFPSLKLLDYH). Scale bar is 1um**

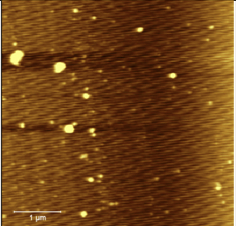

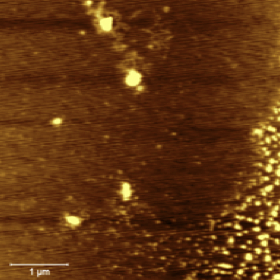

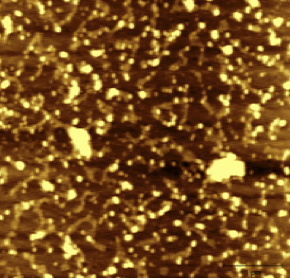

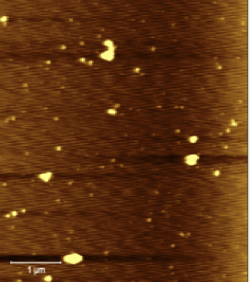

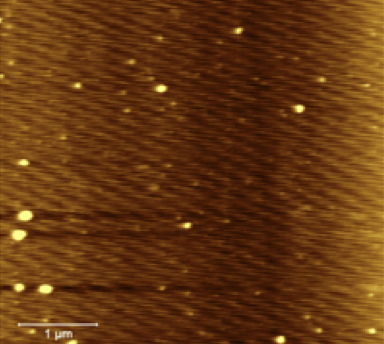

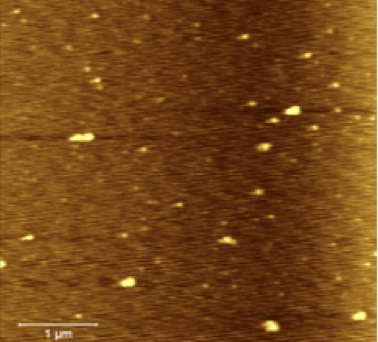

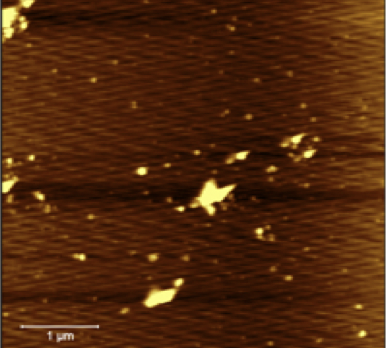

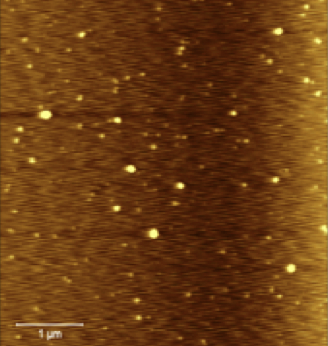

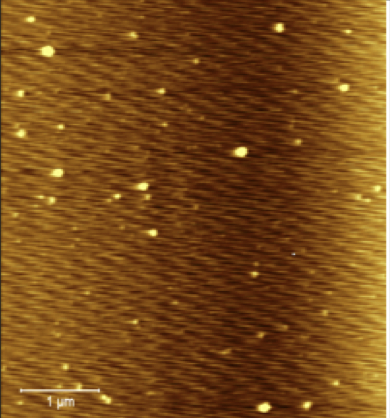

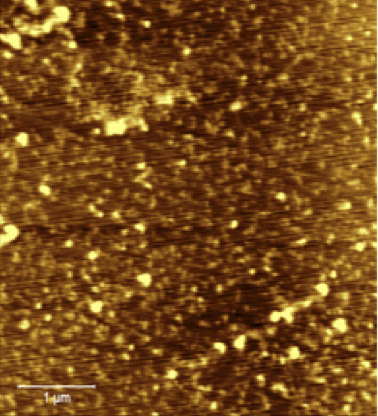

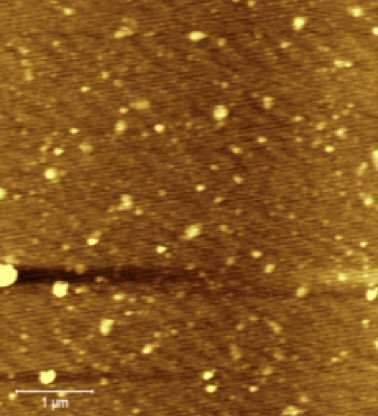

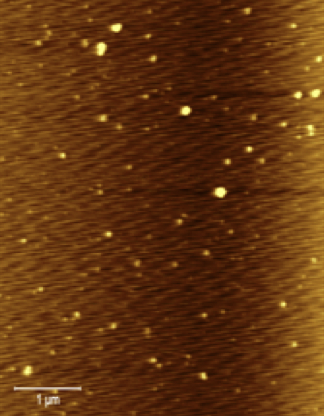

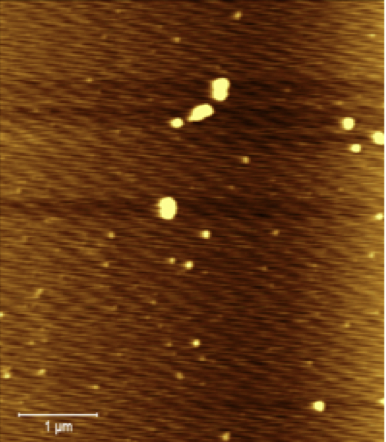

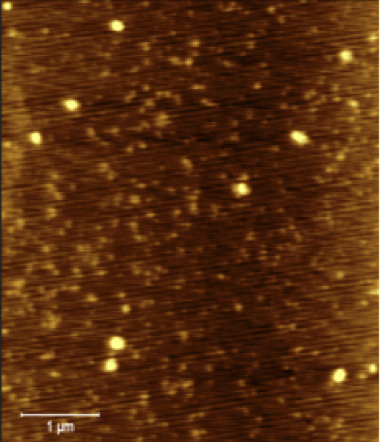

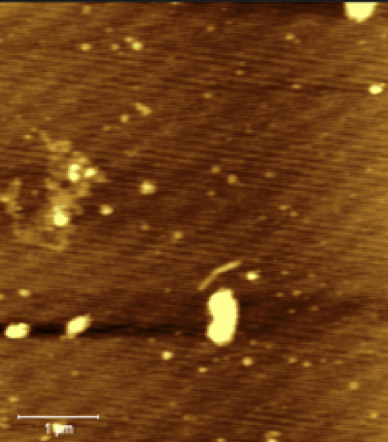

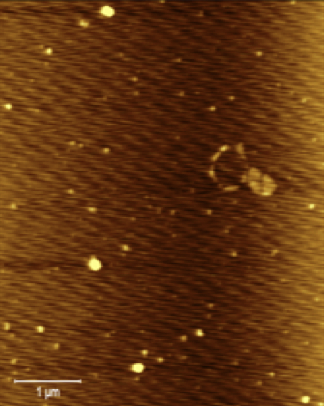

Htt (Q25) Protein

Protein

P1

P3

P5

T= 4

T=0

T=8

T=24

Figure S5. Shows the aggregation kinetics of HttQ25, both in the absence and presence of inhibitory peptides. The AFM images show samples obtained at 0 hour, 4 hours, 8 hours, and 24 hours. The first row of Images corresponds to the fibril growth kinetic of the Htt Q25 protein. The second row of images corresponds to the fibril growth kinetics of HttQ25 in the presence of inhibitory peptide 1 (**HHGANSLLGLVQS).** The third row shows fibril growth kinetics of HttQ25 in the presence of inhibitory peptide 3 (**HGLHSMHNKLLQT)** and the last row shows fibril growth kinetics of HttQ25 in the presence of inhibitory peptide 5 (**WMFPSLKLLDYH). Scale bar is 1um.**

| **Htt Protein**  **Inhibitory**  **Peptide** | **Htt(46Q)** | **Htt (25Q)** | **Htt(103Q)** |
| --- | --- | --- | --- |
| Peptide 1. HHGANSLSLVSQD | Decreased fluorescence | Decreased fluorescence | Decreased  fluorescence |
| Peptide 2. SWPLTPVRFMTE | Decreased fluorescence | Increased  fluorescence | Decreased fluorescence |
| Peptide 3. HGLHSMHNKLTR | Decreased fluorescence | Decreased fluorescence | Decreased fluorescence |
| Peptide 4. FKQDAWEAVDIR | Decreased fluorescence | Increased  fluorescence | Increased  fluorescence |
| Peptide 5. WMFPSLKLLDYH | Decreased fluorescence | Decreased fluorescence | Decreased fluorescence |
| Peptide 6. HVTFKFQWDRES | Increased fluorescence | Decreased fluorescence | Decreased fluorescence |

Table S3. Summarizes the increase or decrease in fluorescence detected after ThT assay for all three Htt proteins, in the presence of the 6 peptides.
